# Recombinant expression, purification, and antifungal activity of the novel antimicrobial peptide TaW662

**DOI:** 10.1101/2024.04.29.591476

**Authors:** Chen Wang, Baoyi Yang, Yanlin Yang, Lu Kang, Fujiao Liu, Zhiwen Wang, Deshun Feng

**Author notes:** **Corresponding author** *Correspondence author:* (D. Feng). Those authors contributed equally to this work (Chen Wang, Baoyi Yang).

## Abstract

*Blumeria graminis* f. sp. *tritici* (*Bgt*) is a significant wheat fungal pathogen, posing threats to both yield and quality. Antimicrobial peptides, with their broad-spectrum activities, hold promise in combating *Bgt*-induced wheat fungal diseases.In this study, we identified *TaW662*, an antifungal peptide gene sourced from the wheat-*Thinopyrum intermedium* disomic alien addition line SN6306. Through third-generation transcriptome sequencing, we obtained the full-length transcript of *TaW662*. Notably, *TaW662* exhibited upregulated expression in response to powdery mildew infection in SN6306. Subcellular localization analysis revealed TaW662’s extracellular secretion, suggesting its role in defense mechanisms. Additionally, the TaW662 protein was expressed in *Escherichia coli*, and the purified protein could inhibit the growth of *Bgt in vitro*. Utilizing the online alphafold2 server, we predicted the three-dimensional structure of TaW662, aiding in understanding its fungicidal mechanisms. Analysis of TaW662’s physicochemical properties further supported its potential efficacy as a fungicide against *Bgt*. In conclusion, TaW662 emerges as a promising candidate for combating *Bgt*-induced wheat fungal diseases, warranting further exploration for agricultural disease management strategies.

**Highlights:** *TaW662*, a secreted protein homologous to *TaWIR1*, is induced in wheat by *Blumeria graminis* f. sp. *tritici* (*Bgt*).

The expression pattern of *TaW662* in wheat under induced by *Bgt* was analysed using RNA-Seq technology.

The three-dimensional structure of TaW662 was predicted using AlphaFold2.

The growth of *Bgt* is inhibited by recombinant TaW662.

## 1. Introduction

The global increase in population underscores the urgent need for enhanced food production, with wheat standing as a crucial staple crop worldwide. However, fungal diseases severely compromise both the yield and quality of wheat. The prevalent use of chemical fungicides poses risks to microbial communities and undermines efforts to maintain ecological balance [1]. Consequently, there is an acute demand for low-pollution and highly efficient biological fungicides. Plant antimicrobial peptides (AMPs) emerge as promising candidates for this purpose, owing to their natural origins, potent antimicrobial properties, and broad spectrum of resistance.

AMPs are natural molecules characterized by their small molecular weight, positive charge, and amphiphilic properties [2,3]. They possess the ability to inhibit various pathogens, including bacteria and fungi, without inducing pathogen resistance, making them viable alternatives to chemical fungicides [4].

Plant AMP genes are abundant and expressed in various plant parts such as roots, leaves, flowers, fruits, and seeds, contributing significantly to resistance against pathogen infections [5,6]. AMPs are classified based on their amino acid composition into cysteine-rich AMPs (CrAMPs), proline-rich AMPs (PrAMPs), and glycine-rich AMPs. Most plant antimicrobial peptides are rich in cysteine residues, forming secondary structures known as cysteine-stabilized α-β motifs [7].

CrAMPs typically exert their antifungal activity by interacting with cell membranes. For instance, Dahlia merckii antimicrobial protein 1 exhibits its mode of action by binding to sphingolipids on the cell membrane of *Saccharomyces cerevisiae* [8]. Additionally, the synergistic effect of AMPs with compounds like jasmonic acid and glycerol-3-phosphate can enhance plant resistance [9].

In this study, a differentially expressed gene, TaW662, was identified from the suppression subtractive hybridization (SSH) library of the wheat-*Thinopyrum intermedium* disomic alien addition line SN6306. Sequence analysis revealed a high homology between TaW662 and TaWIR1. TaWIR1, initially discovered in wheat [10], has subsequently been identified in barley and rice as well [11,12]. WIR1 expression is induced by both *Blumeria graminis* f. sp. *tritici* (*Bgt*) and *Puccinia striiformis* f. sp. *tritici* in wheat [12,13]. In rice, WIR1 homologs have been found to enhance resistance against *Magnaporthe grisea* [12]. Notably, TaWIR1 is characterized by an abundance of glycine and proline residues, potentially facilitating adhesion between the plasma membrane and the cell wall [14]. Tufan et al. have suggested that the highly conserved sequence of TaWIR1 protein plays a pivotal role in plant defense against pathogen infections [15].

*TaW662*, identified as a homolog of *TaWIR1*, exhibits significant induction in response to *Bgt*, indicating its role as an antifungal peptide resistant to *Bgt*. This was confirmed through cloning, prokaryotic expression, and subsequent antimicrobial activity analysis.

## 2 Material and methods

### 2.1 Plants, vectors, strains, and growth medium

The materials utilized in this study comprised the wheat-*Thinopyrum intermedium* disomic alien addition line SN6306, wheat YN15, and Race E09 of *Bgt*. Further information regarding these materials is available in Supplementary Material 1.

### 2.2 Gene cloning and bioinformatic analysis

The SSH library of SN6306 induced by *Bgt* had been previously constructed. Subsequently, based on one of the differentially expressed genes, a pair of primers (Forward: 5 ′ -CATCGAATCATCAGCATGGC-3 ′, Reverse: 5 ′ -ACGGGTGTAATTAGGGCTGG-3 ′) was designed to amplify the novel gene *TaW662* from SN6306’s genomic DNA and complementary DNA (cDNA) through PCR. Gene structure and protein sequence comparison are provided in Supplementary Material 2.

### 2.3 Subcellular localization analysis

The TaW662-GFP fusion expression vector was constructed using the pCambia 13000221-GFP.3 vector as the plasmid and Agrobacterium GV3101 as the strain. Detailed protocols for the construction of the expression vector and subsequent subcellular localization experiments are provided in Supplementary Material 3.

### 2.4 Expression analysis of the *TaW662* gene induced by *Bgt* using the transcriptome

The transcript expression changes of *TaW662* in the epidermal cells of SN6306 leaves induced by *Bgt* were analysed. One week old leaves were inoculated with powdery mildew spores *E09* and subsequently cultured [16]. Epidermal cells were harvested from 40 wheat leaves, both inoculated and non-inoculated, at 12, 24, and 48 hours post-inoculation. Three biological replicates were collected at each time point for second-generation transcriptome sequencing, while a pooled sample from each time point was subjected to PacBio third-generation full-length transcriptome sequencing. Sequencing service was provided by Personalbio Biotechnology Co., Ltd. Shanghai, China.

### 2.5 Construction of prokaryotic expression vector

A pair of specific primers was designed for cloning TaW662. According to SignalP-5.0 prediction, the signal peptide cleavage site of TaW662 was identified between amino-acid residues 32nd and 33rd. Subsequently, a TaW662 gene fragment without the signal peptide and optimized for *Escherichia coli* codon preference was synthesized. This fragment was then inserted into the expression vector pET32a between the *Nco* I and *EcoR* I sites. The synthesis and insertion steps were carried out by Boshan Biotechnology Co. Ltd. The resulting prokaryotic expression construct, pET32a-TaW662, contained His-tag and enterokinase cutting sites before the insertion sequence. Following transformation into *E. coli* BL21 (DE3) plysS, positive transformants were confirmed through nucleotide sequencing.

### 2.6 Expression of the fusion protein

The recombinant plasmid was introduced into the host strain BL21 (DE3) plysS. Single colonies were selected and inoculated into LB liquid medium supplemented with ampicillin antibiotics (100 μg/mL), followed by cultivation at 37°C for 16 hours. Positive recombinants, confirmed by sequencing, were then cultured in LB liquid medium. For protein expression, the positive recombinants were cultured in autoinduction medium supplemented with 100 μg/mL ampicillin and containing glucose, lactose, and glycerol as carbon sources, preferred by *E. coli*, in a volume ratio of 5%. Induction was carried out in an oscillator under autoinduction conditions (200 rpm at 37°C for 5 h and 200 rpm at 25°C for 16 h). Subsequently, protein expression was assessed using 12% SDS-PAGE electrophoresis. The TaW662 protein was detected through Western blot analysis.

### 2.7 Antifungal assay

After the purification and refolding of the prokaryotically expressed proteins, they were applied by spraying onto the leaves of the wheat cultivar YN15. Subsequently, the leaves were inoculated with *Bgt*, with the His-tagged protein serving as a control. The growth of *Bgt* spores was then observed and photographed under a microscope to analyse the influence of TaW662 on powdery mildew infection. To quantify the extent of infection, Image J software was utilized to analyse the pixels of the *Bgt* infection area (BIA) and normal leaf area (NLA). The ratio of the pixels of the *Bgt* infection area to the total pixels of the leaves was calculated as previously described [17]. Further details are provided in Supplementary Material 4.

## 3 Results

### 3.1 Sequence and structure analysis of TaW662

The gene *TaW662*, identified as differentially expressed in the SSH library, was found to be located on chromosome 5B of Chinese Spring. The *TaW662* gene comprises 373 base pairs, encoding a polypeptide of 81 amino acids. An intron is situated between bases 124 and 250. (Fig. 1A). Alignment of the amino acid sequences of TaW662 in NCBI revealed 10 homologous proteins from *Triticum aestivum, Triticum durum, Hordeum vulgare, Aegilops tauschii*, and *Triticum urartu*. Comparing TaW662 with these homologous proteins using DNAMAN software showed a sequence similarity of 70% (Fig. 1B). Notably, among the 10 homologous proteins, five were identified as WIR proteins, while the remaining five were not annotated.

**Fig. 1.**
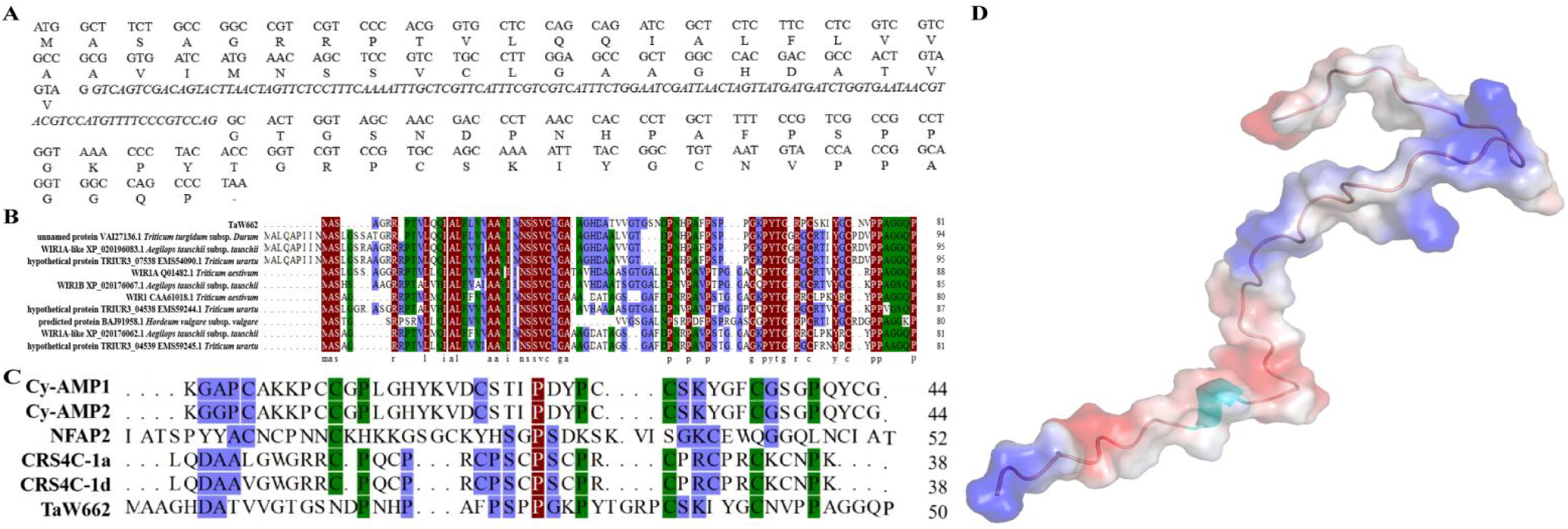
TaW662 sequence analysis and 3D structure prediction. (A) *TaW662* has two exons and one intron. The intron is shown in italics and is located between bases 124 and 251. (B) TaW662 and 10 homologous proteins were compared using DNAMAN software. (C) Multisequence alignment of AMPs. (D) TaW662 3D structure prediction model based on coverage and pLDDT score.

As a result of its high sequence identity with known antifungal peptides, TaW662 is proposed to be an antimicrobial peptide (AMP). To validate this, TaW662 was submitted to the APD database for comparison with the AMPs in the database. The results revealed that TaW662 shares a sequence identity of 41.19% with the five AMPs (Fig. 1C). Further analysis of the structural properties of TaW662 and the five AMPs is provided in Supplementary Table 1. Notably, the pI (isoelectric point) of TaW662 was determined to be 8.08, with a hydrophobic residue ratio of 24% and a positive Boman index. These predicted values closely resemble those of the other five AMPs. Additionally, the 3D structure of TaW662 was predicted and analysed using alphafold2 (Fig. 1D). The optimal structural model of TaW662 was constructed based on the coverage of each model and the pLDDT score assigned to every amino acid, and reconstructed the model based on protein surface potential and secondary structure (Supplementary Figure. 1).

### 3.2 TaW662 can be secreted outside the cell and the gene is up-regulated by powdery mildew

We transiently expressed TaW662-GFP under the control of the 35S promoter in *N. benthamiana* leaves, with GFP serving as the control. After 2 days of culturing, we analysed the subcellular localization using confocal laser scanning microscopy under a 488 nm excitation wavelength. The results demonstrated that TaW662-GFP was observed to be secreted outside the cell (Fig. 2A). These findings suggest that TaW662 may undergo synthesis and secretion into the intercellular space to exert its functional role.

**Fig. 2.**
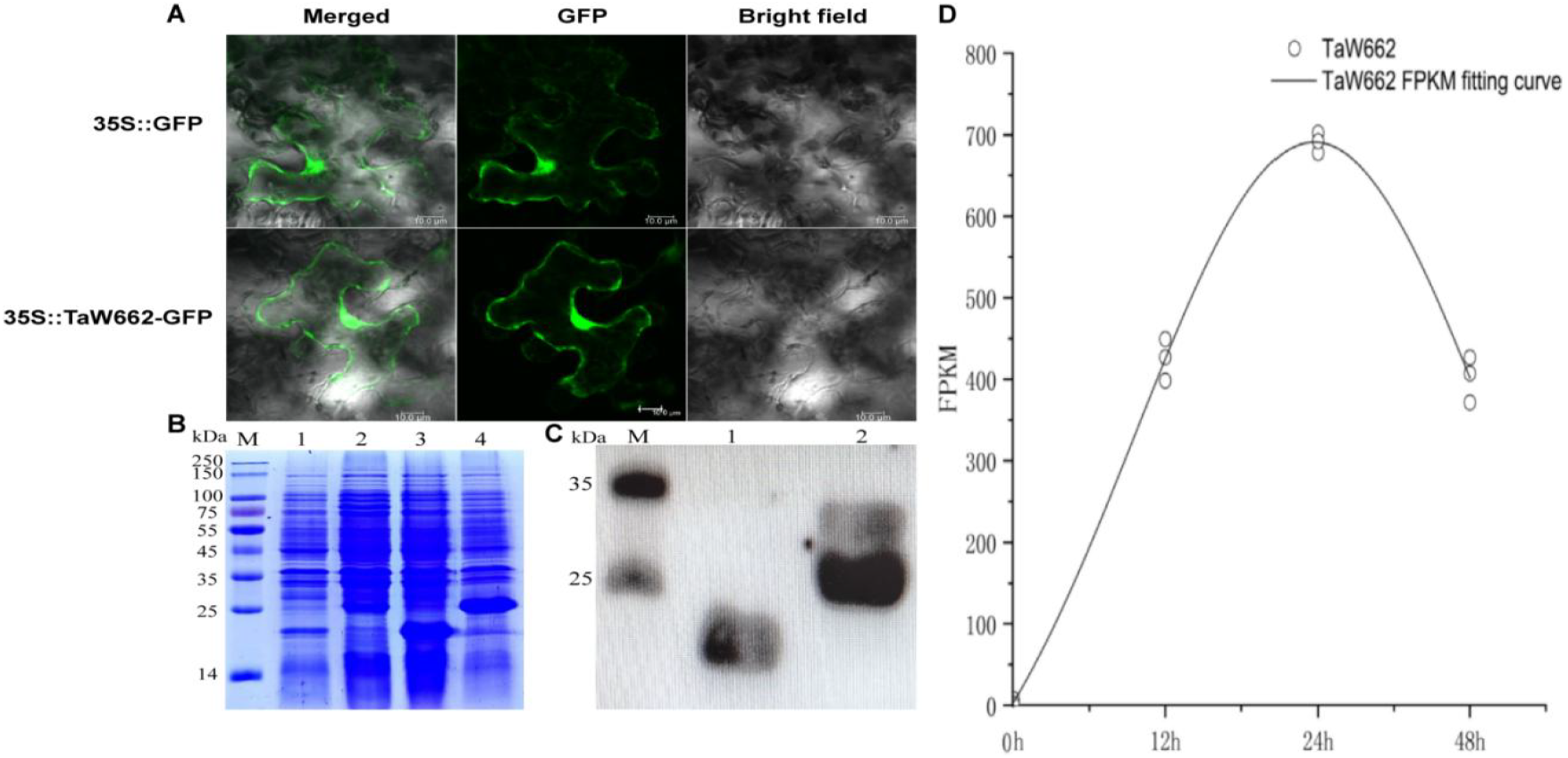
TaW662: Subcellular Localization, Prokaryotic Expression, and Expression Analysis. (A) Subcellular localization of the fusion protein in *N. benthamiana* leaf epidermal cells. The constructs *35S*::*TaW662*-*GFP* and *35S*::*GFP* (control) were infiltrated into *N. benthamiana* leaf cells via *Agrobacterium*-mediated transformation. Green fluorescent protein (GFP) fluorescence was visualized using confocal laser scanning microscopy. Scale bar = 10 μm. (B) Detection of protein expression using 12% SDS-PAGE. M: Broad molecular weight protein ladder; Lane 1: Uninduced pET32a crude protein; Lane 2: Uninduced TaW662 crude protein; Lane 3: Induced pET32a crude protein; Lane 4: Induced TaW662 crude protein. (C) Expression of the fusion protein detected by Western blotting. M: Blue Plus®II Protein Marker; Lane 1: Protein of pET32a; Lane 2: Protein of TaW662. (D) Expression level of *TaW662* calculated using the FPKM method. The y-axis represents the gene expression value (FPKM), and the x-axis denotes sequential developmental time. Each point represents a single sample, with black lines indicating the trends in the average level of gene expression.

The fusion protein containing TaW662 was successfully produced through prokaryotic expression of the pET32a-TaW662 construct. The recombinant protein exhibited a molecular weight of approximately 25.0 kDa, whereas the empty vector (pET32a) displayed a molecular weight of around 20.0 kDa (Fig. 2B). The presence of the His-tag protein was confirmed on the PVDF membrane via Western blot analysis using the anti-His-tag antibody (Fig. 2C).

The expression levels of *TaW662* gene in leaves epidermal cells of SN6306 were determined using the Htseq software and the FPKM method. The control represented the expression level of TaW662 without inoculation. Notably, following inoculation for 12, 24, and 48 hours, the expression level of TaW662 gradually increased and reached 690 on the second day of inoculation (Fig. 2D).

### 3.3 TaW662 inhibits *Bgt* infection

To validate the function of the TaW662 protein, we inoculated *Bgt* on YN15 wheat leaves sprayed with TaW662 and a control tag protein respectively. After 3 days of inoculation, the infection stages of spores were counted. At least 200 spores were counted on each wheat leaf to observe their life cycle, and three leaves were used in every treatment (Fig. 3A). The results indicated a significant inhibition of spore growth by TaW662 (Fig. 3B). After 5 days of inoculation, the infection ratio on the leaves of the control group reached 11.35%, whereas the experimental group treated with TaW662 had fewer spores, resulting in an infection ratio of only 2.07% (Fig. 3C). The haustorial index (HI) was calculated as the ratio of haustoria-containing infected cells, with the control group having a HI of 59.2%. Comparatively, the TaW662-treated group exhibited a HI of only 42.23% after 3 days of inoculation, indicating reduced infection (Fig. 3D).

**Fig. 3:**
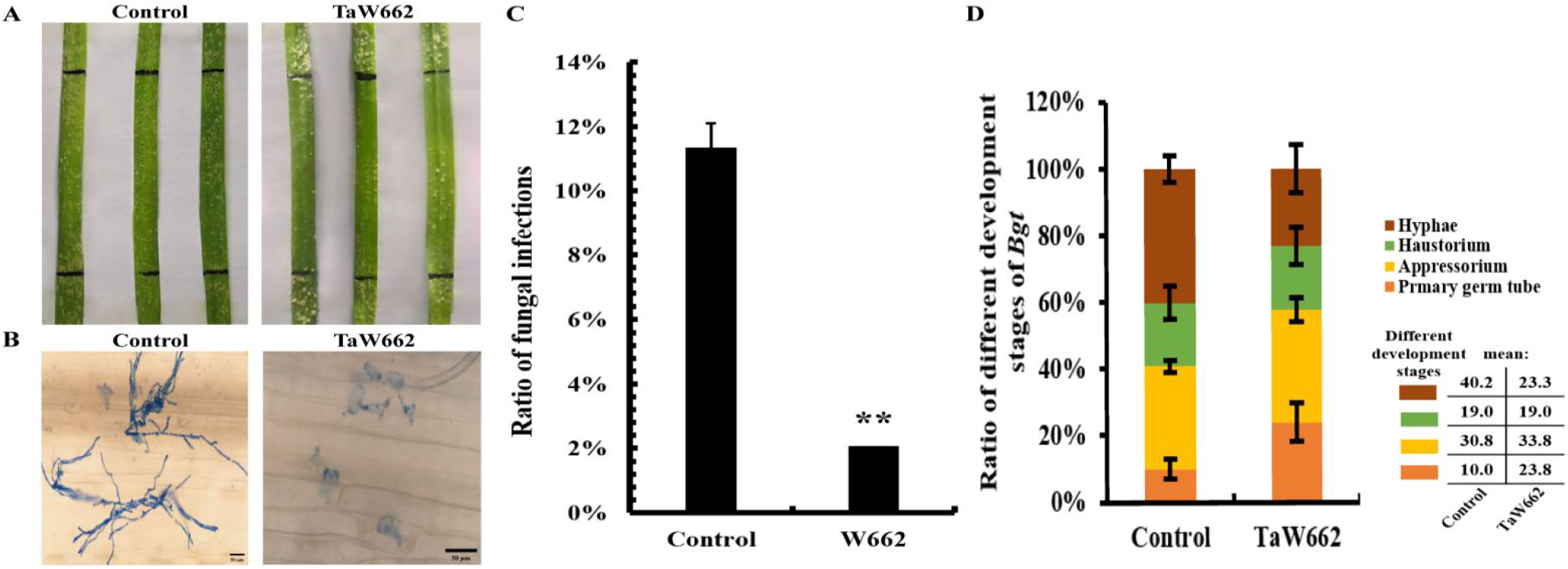
Fungal inhibition results of leaves sprayed with TaW662. (A) The leaves of wheat SN6306 were sprayed with TaW662 protein and control (CK), then inoculated with E09 for 5 days. Black lines indicate the areas (5 cm) sprayed with protein on the leaves. (B) Image of *Bgt* spores after 3 days post-inoculation on wheat SN6306 leaf epidermal cells. The scale bar in the control image is 50 μm. (C) Ratio of *Bgt* infection area on leaves. (D) Ratio of different calculated stages of *Bgt* spore growth after 3 days of inoculation. Ratio of infections =BIA/NLA. Haustorial index (HI) was calculated as the ratio of haustoria-containing infected cells. HI = (Hypha + Haustorium)/ Infected cells

## 4 Discussion

Fungal diseases pose a significant threat to crop yield and quality, yet effective control methods remain limited. Chemical fungicides are commonly employed for disease management, albeit with potential negative impacts on microbial community richness and diversity within the ecological environment [1]. Antimicrobial peptides (AMPs) offer a promising alternative, as they effectively inhibit pathogen activity across a broad spectrum without promoting drug resistance. Notably, various types of AMPs have been identified in plants [18], including antifungal genes discovered in wheat crops [19,20].

In our previous study, we constructed a SSH library from the SN6306 infected by *Bgt*. Through screening, we identified several sequences associated with disease resistance and defense mechanisms. Among them, a novel antimicrobial peptide gene named *TaW662* was cloned from SN6306. Sequence alignment using the URGI database revealed that *TaW662* is located on chromosome 5B of Chinese Spring. Transcriptome sequencing and gene expression analysis conducted on leaves of SN6306 demonstrated that the expression of *TaW662* was induced in response to *Bgt* infection.

Among the results of the multiple sequence alignment, TaW662 exhibited the highest similarity to Cy-AMP1 and Cy-AMP2, both of which are rich in proline residues. Their physicochemical properties, including pI values, hydrophobic residue ratios, Boman indexes, positive charge numbers, α-helix prediction values, and β-turn prediction values, were found to be proximate. These indexes are closely associated with the antifungal effects of AMPs [21].

Taniguchi et al. isolated several peptides with high pI values from soybean and rice bran protein hydrolysates and identified their antimicrobial activities [22,23]. Cationic antimicrobial peptides typically possess 2-9 positive charges and contain up to 50% hydrophobic amino acids. Upon interaction with the cell surface, these peptides aggregate and fold into specific secondary structures, facilitating insertion into the cell membrane [24]. Barros et al. reported on a recombinant peptide, Jaburetox-2Ec, which displayed a hydrophobic residue ratio of 34% and formed a β-turn, enabling it to penetrate lipid vesicle membranes and induce membrane permeability [25]. Upon binding of positively charged residues to negatively charged lipids on the membrane, the peptide chain adopts an α-helix structure [26]. Disulfide bonds stabilize the α-helix and form a hydrophobic cavity, facilitating the extension of hydrophobic peptide molecules into the cell membrane and the formation of ion channels [27]. However, both Cy-AMP1 and Cy-AMP2 also possess a high proportion of cysteine residues, which may interact to form cysteine-stabilized α-β motifs, crucial for exerting antifungal effects [28]. The cysteine proportion in TaW662, lacking signal peptides, is lower than in other AMPs, potentially hindering the formation of an α-helix structure. In contrast, proline content accounts for 20% of the total amino acids. Therefore, TaW662 may exert its antifungal activity differently from Cy-AMP1 and Cy-AMP2.

Blast analysis of the sequence alignment revealed 10 proteins with high similarity to TaW662, among which five were known as WIR proteins, while the others were not annotated. These sequences originate from wheat and its wild relatives. Members of the *WIR* gene family can be induced by various pathogens, such as wheat leaf rust, and have been identified from wheat, barley, and *A. tauschii* [29,30]. *TaWIR1* encodes a small protein rich in glycine and proline, which may facilitate cell wall contact [15]. Tufan et al. suggested that TaWIR1 is resistant to *Magnaporthe oryzae* but does not affect wheat *Bgt* [15]. However, our study suggests that the externally applied recombinant protein TaW662 may inhibit *Bgt* invasion.

In the subcellular localization experiment, TaW662-GFP fluorescence was observed in the intercellular space, indicating that TaW662-GFP was secreted outside the plasma membrane. This suggests that TaW662 might play a role in disease resistance and defense outside the cell. Similar localization patterns have been reported for previously studied rice antimicrobial peptides OsDEF7 and OsDEF8, which are also localized in the periphery of cells [31]. This localization pattern of TaW662 may contribute to resisting the early infection of pathogens.

The TaW662 protein was expressed in Escherichia coli by constructing the expression vector pET32a-TaW662. The recombinant protein was expressed in the form of soluble protein through autoinduction. To verify the antifungal activity of TaW662 in vitro, wheat leaves were sprayed with the TaW662 protein and then infected with *Blumeria graminis* f. sp. *tritici* (*Bgt*). Before the experiment, the tag sequence of the recombinant protein was removed by enterokinase digestion to ensure that the TaW662 protein was responsible for the antifungal activity. In the early stages of infection, TaW662 inhibited the development of *Bgt* spores, causing them to remain in the primary germ tube and appressorium stage, thus delaying spore growth.

## 5 Conclusion

The TaW662 protein, sharing homology with TaWIR, exhibits inducibility by powdery mildew and is secreted into the intercellular space to combat powdery mildew invasion.

## CRedit authorship contribution statement

**Chen Wang**: Writing - original draft, Visualization, Methodology, Formal analysis, Investigation. **Baoyi Yang**: Writing - original draft, Visualization, Methodology, Formal analysis, Investigation. **Yanlin Yang**: Visualization, Validation, Investigation. **Lu Kang**: Visualization, Investigation. **Fujiao Liu**: Visualization, Investigation. **Zhiwen Wang**: Visualization, Investigation. **Deshun Feng**: Writing - review & editing, Project administration, Methology, Supervision, Funding acquisition, Conceptualization.

## Abbreviations

AMP: antimicrobial peptide
SSH: suppression subtractive hybridization
GFP: green fluorescent protein
FPKM: Fragments Per Kilo bases per Million fragments
SN6306: Wheat varieties Shan Nong 6306
YN15: Wheat varieties Yan Nong 15
WIR: wheat-induced resistance

---

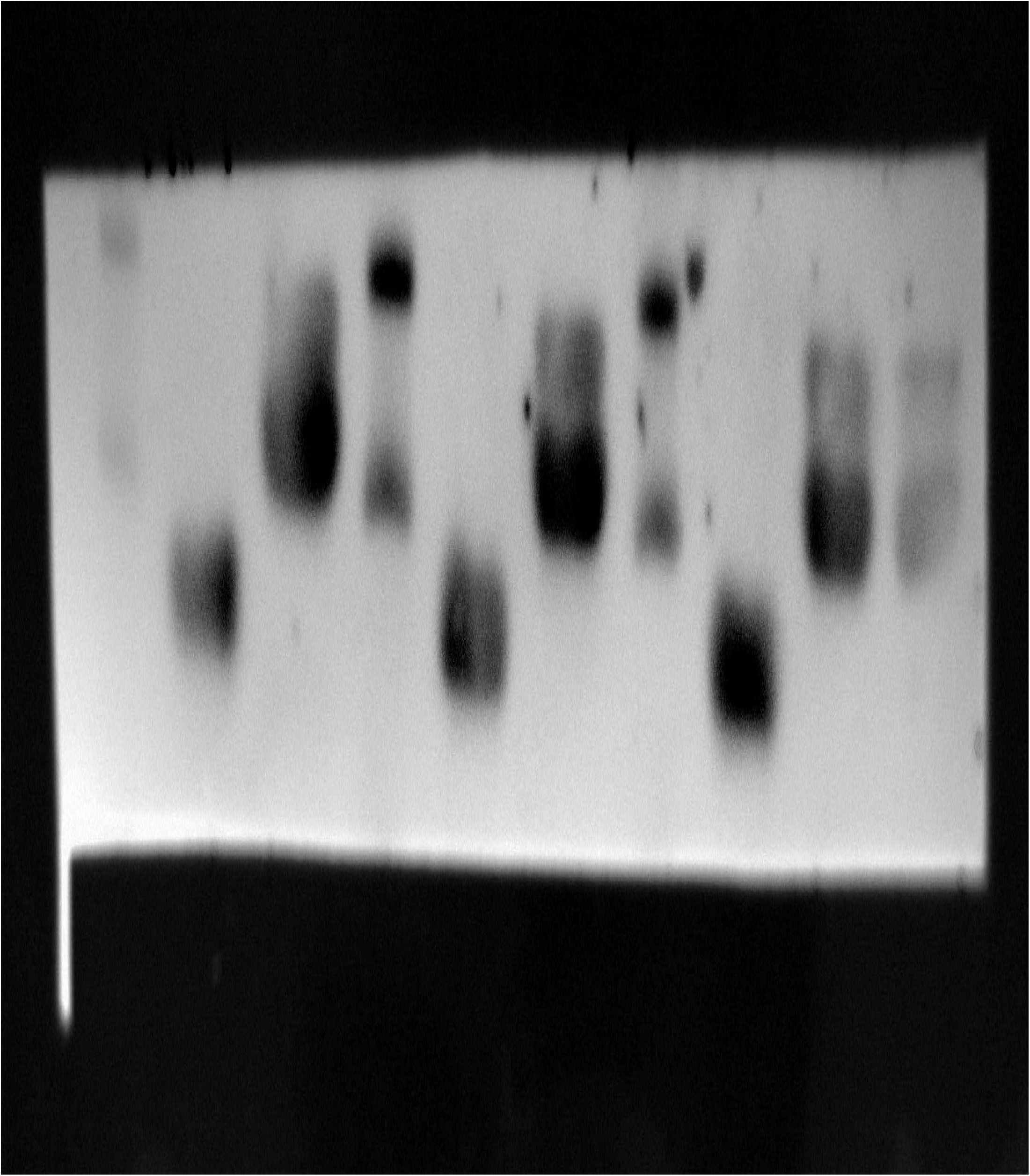

