## Supplemental Files for "Recombinant expression, purification, and antifungal activity of the novel antimicrobial peptide TaW662"

**Supplemental Material**

**1 Plants, vectors, strains, and growth medium**

The wheat cultivar Yannong 15 (YN15), which is highly susceptible to *Bgt*, was cultured in an incubator under light conditions for 16 h at 20°C and under dark conditions for 8 h at 18°C. The transcriptome data of the wheat-*Thinopyrum intermedium* additional line SN6306 with high resistance to *B. graminis* f. sp. *tritici* were obtained by sequencing [1].

Race *E09* of *Blumeria graminis* f. sp. *tritici* was subcultured in an incubator at 20°C. The expression vector pET32a was stored at -20°C. *Escherichia coli* DH5α competent cells and *E. coli* BL21 (DE3) plysS competent cells were purchased from TransGen Biotechnology Co. Ltd. and then stored at -80°C.

**2 Gene cloning and bioinformatic analysis**

Sequence alignment was performed using the URGI database. The gene structure of *TaW662* was predicted using Softberry and signal peptide site prediction was performed using SignalP-5.0, and its 3D structure was modeled using alphafold2 and Pymol. Protein sequence comparison was performed using DNAMAN. In addition, the antimicrobial peptide properties of TaW662 were calculated and predicted using APD3.

URGI: https://wheat-urgi.versailles.inra.fr/

Softberry: http://www.softberry.com/berry.phtml?topic=fgenesh&group=programs&subgroup=gfind

SignalP-5.0: https://services.healthtech.dtu.dk/services/SignalP-5.0/

Protein Multiple Sequence Alignment: DNAMAN

APD3: https://aps.unmc.edu/AP/

Alphafold2: https://colab.research.google.com/github/sokrypton/ColabFold/blob/main/AlphaFold2.ipynb.

**3 Subcellular localization analysis**

The TaW662 green fluorescent protein (GFP) fusion expression vector was constructed. The cDNA fragment of *TaW662* was inserted between the *Xba* I and *Asc* I sites of the pCambia13000221-GFP.3 vector and then transformed into *Agrobacterium* GV3101. The positive strains were cultured in LB liquid medium containing 20 μM Acetosyringone (AS), 50 mg/L kanamycin, and 100 mg/L rifampicin and grown overnight at 28°C with shaking at 200 rpm. The cultures harvested by centrifugation were resuspended in a solution containing 10 mM MgCl_2_, 0.1 mM AS, and 10 mM 2-morpholinoethanesulfonic acid with pH 5.2 and an OD_600_ of 0.6. The cultures were injected into the lower epidermis of 6-8 leaves of *Nicotiana benthamiana* by using a 1 mL syringe with the needle removed. The injected plants were left in the dark for 12 h and then incubated under light conditions for 16 h at 21°C and under dark conditions for 8 h at 18°C. Plasmolysis was performed with 0.8 M mannitol solution using the same method as *Agrobacterium* GV3101. Subsequently, leaf discs were immediately excised using a punch for observation, the treated leaves were sampled and observed under a confocal laser scanning microscope at an excitation wavelength of 488 nm. The leaves injected with pCambia 1300221-GFP.3 were used as control.

**4 Antifungal assay**

Wheat YN15 was cultured for 5 days, leaves were fixed on an acrylic plate with rubber bands to retain all leaves on a horizontal plane, and a 5 cm width was left in the middle of the rubber bands. Purified TaW662 protein was digested with enterokinase and mixed with Tween 20 (*V*/*V*: 0.025%). The enzyme digestion was performed as follows. Every 300 μL (concentration: 241-252 μg/mL) of purified protein was added with 0.06 U recombinant enterokinase and reacted in a metal bath at 25°C for 16 h. The protein solution was sprayed on the leaves fixed by a rubber band with a cotton swab. After the solution dried, the spores of *Bgt* were sprayed onto the treated wheat leaves and then placed in an incubator at 25°C for 16 h light/day. After 3 days, the leaves were harvested, soaked in the decolorizing solution overnight, bleached with sterile water twice, and then dyed in dyeing solution for 15-30 min. After washing three times with sterile water, spore development can be observed under the microscope. Leaves sprayed with pET32a crude tag protein were used as control. After 6 days of inoculation, the remaining treated leaves were harvested. Images of the wheat leaves were taken to analyze the infection ratio of *Bgt*. Image J software was used to analyze the pixels of *Bgt* infection area and normal leaf area, and the ratio of the pixels of the *Bgt* infection area to the total pixels of the leaves was calculated as previously described by Karisto *et al.* [2]. The ratio was used as the area ratio of fungal infection.

**Supplementary Table**

**Supplementary Table 1 TaW662 AMP comparison results**

| **Parameter** | **Source** | **Number of amino acids** | **Molecular weight (kDa)** | **Theoretical pI** | **Hydrophobic residue（%）** | **Boman index（kcal/mol）** | **Total net charges** | **α-helix（%）** | **β-turn（%）** |
| --- | --- | --- | --- | --- | --- | --- | --- | --- | --- |
| **Tools** | NCBI/URGI | DNAMAN | DNAMAN | ProtParam | APD | APD | APD | SOPMA | SOPMA |
| **TaW662** | *Wheat* | 49 | 4.84 | 8.08 | 24 | 1.16 | +1.5 | 0 | 8.16 |
| **Cy-AMP1** | *Cycas revoluta* | 44 | 4.59 | 8.47 | 31 | 0.71 | +3 | 0 | 6.82 |
| **Cy-AMP2** | *Cycas revoluta* | 44 | 4.58 | 8.47 | 29 | 0.73 | +3 | 0 | 6.82 |
| **NFAP2** | *Neosartorya fischeri* | 52 | 5.56 | 9.02 | 28 | 1.58 | +7 | 0 | 5.77 |
| **CRS4C-1a** | *Mus musculus* | 38 | 4.25 | 9.24 | 36 | 2.75 | +7 | 2.63 | 15.79 |
| **CRS4C-1d** | *Mus musculus* | 38 | 4.23 | 9.24 | 36 | 2.77 | +7 | 2.63 | 10.53 |

**Supplementary Figure**

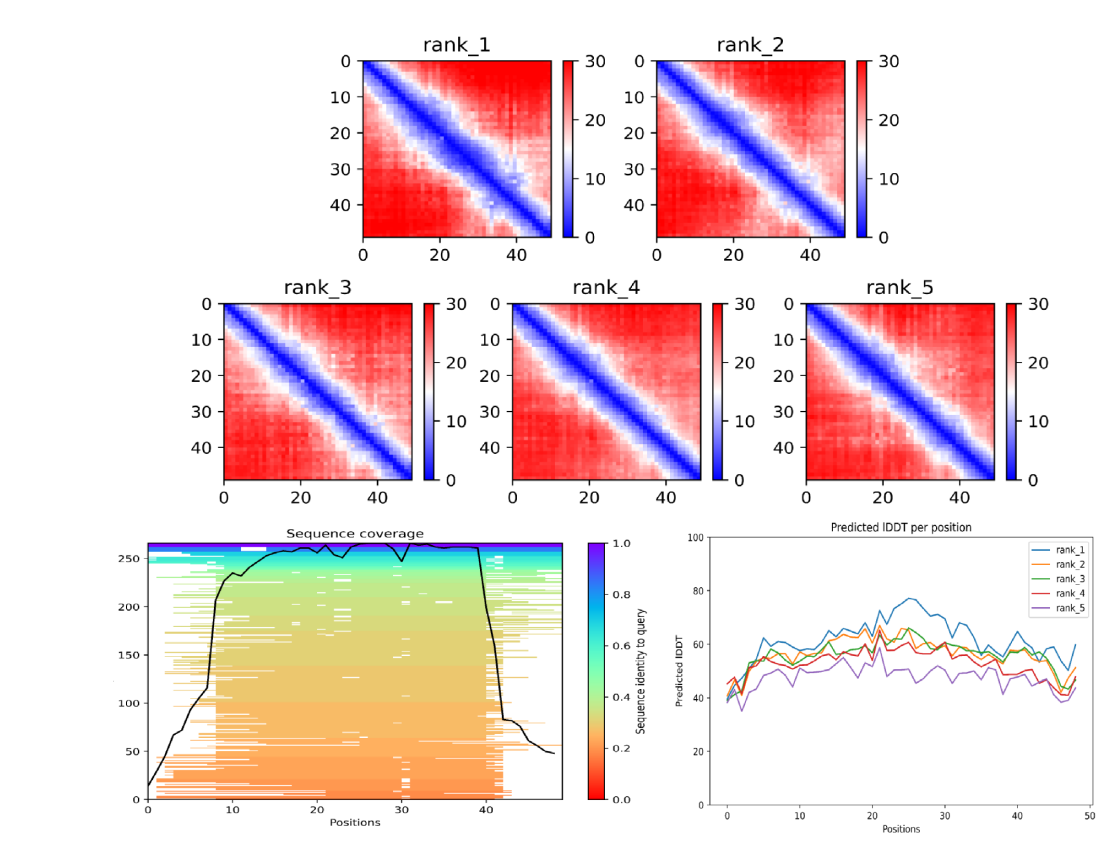

**Supplementary Figure 1 Information about each model predicted by Alphafold2.** Basic parameters of the prediction model based on alphafold2, showing that the rank_1 model is the optimal model in both coverage and pLDDT score.
